# Impacts of high pathogenicity avian influenza H5N1 2.3.4.4b south of the Antarctic Circle

**DOI:** 10.1101/2025.04.13.648652

**Authors:** Simon B.Z. Gorta, Victor Neira, Michelle Wille, David Gajardo, Atilla Yılmaz, Özgün Oktar, Burcu Özsoy, Çağan H. Şekercioğlu

**Author notes:** **Author contributions:** SBZG and ÇHŞ conceived the study; SBZG led the field work; SBZG, MW, AY, ÖO, BÖ and ÇHŞ contributed to field sampling; VN and DJ performed the virus sample analyses; SBZG wrote the first version of the manuscript; VN, MW, DG, AY and ÇHŞ provided inputs.

## Abstract

High pathogenicity avian influenza (HPAI) H5N1 2.3.4.4b poses a substantial conservation threat to ecosystems, populations, and species globally, with its continued spread into new regions increasing concern for potential ecological consequences. During surveys in February – March 2025, we confirmed the virus’s presence at the southern extent of its known range along the Western Antarctic Peninsula, with recorded mortalities in South Polar Skuas S*tercorarius maccormicki* on distinct islands in Marguerite Bay, as well as one confirmed and one suspected case in Kelp Gulls *Larus dominicanus*. At the time of sampling, no evidence of infection was observed in other seabird or mammal species. Consistent with previous global reports, skuas – here, South Polar Skuas – appear particularly vulnerable, yet broader impacts on the local seabird and mammal community remain unclear. Additionally, our use of rapid antigen tests (VDRG® AIV Ag Rapid kit 2.0 – Median Diagnostics) in the field demonstrated their potential utility for real-time surveillance, though false negatives (10%) highlight limitations in test sensitivity. These findings contribute to a growing understanding of the impacts of HPAI H5N1 2.3.4.4b outbreaks on Antarctic species and populations, and will inform continued monitoring, conservation strategies, and biosecurity measures in response to the virus’s ongoing spread.

## INTRODUCTION

High pathogenicity avian influenza (HPAI) H5N1 2.3.4.4b has caused mass mortalities of wild birds and mammals, resulting in a global panzootic since 2021 (Klaassen and Wille 2023; Peacock et al. 2025). The near-global spread of the disease, its capacity to infect and spread amongst many avian and mammalian species, and the scale of mortality are unprecedented in the context of prior HPAI outbreaks (Klaassen and Wille 2023; Peacock et al. 2025). Seabirds and marine mammals have been particularly severely impacted, with many experiencing mass mortality events and some subject to globally significant population declines (e.g., Uhart et al. 2024; Tremlett et al. 2024; Pardo-Roa et al., 2025). As such, the recent spread of the virus into the sub-Antarctic Atlantic and Indian oceans, and the Antarctic Peninsula, which are significant breeding and foraging grounds for many seabirds and marine mammals, is of conservation concern (Banyard et al. 2024; Clessin et al. 2025).

HPAI H5N1 2.3.4.4b was first detected in the sub-Antarctic region in October 2023, and on the Antarctic Peninsula in 2024, with spread along the Antarctic Peninsula as far south as James Ross Island (63°48’ S 57°49’ W) in March 2024 (Bennett-Laso et al. 2024; Kuiken et al. 2025). In the austral summer of 2024/25, the virus continued to cause outbreaks along the Antarctic Peninsula, with detections south of the Antarctic Circle (León et al. 2025; https://scar.org/library-data/avian-flu). In addition to continued amplification along the peninsula, a substantial spread event, to the sub-Antarctic Islands of Marion, Crozet and Kerguelen occurred (Clessin et al. 2025). While many seabird and marine mammal populations are abundant throughout the regions where these outbreaks have occurred, only some species appear to be impacted by the virus. Notable amongst these are skuas (family *Stercorariidae*) which exhibit a variety of foraging and social behaviours such as predation, kleptoparasitism, scavenging, and the use of communal freshwater bathing and drinking sites, which are likely to readily facilitate viral spread (Camphuysen et al. 2022; Gorta et al. 2024).

Globally, HPAI H5N1 2.3.4.4b mortalities have been detected in six of the seven skua species, including a decline of roughly 50% of the global population of Great Skua *Stercorarius skua* in the northern hemisphere in 2021/2022 (Gorta et al. 2024; Tremlett et al. 2024). In the subantarctic and Antarctic regions, mortality events have occurred in both Brown *Stercorarius antarcticus* and South Polar Skua *Stercorarius maccormicki* (Banyard et al. 2024; Bennett-Laso et al. 2024; León et al. 2025). In contrast, few mortalities have yet been reported in penguin species, shags, or even marine mammals along the Antarctic Peninsula.

In this study, we conducted comprehensive surveys of live and dead seabirds and marine mammals on Horseshoe and Dismal Islands in Marguerite Bay in the southern Western Antarctic Peninsula. To date, most sampling efforts have largely occurred in northern regions on or adjacent to the Antarctic Peninsula, with limited sampling south of the Antarctic Circle (Banyard et al. 2024; Bennett-Laso et al. 2024; but see León et al. 2025). These sites are highly remote, and to our knowledge, Dismal Island represents the furthest southern location sampled for HPAI H5N1 2.3.4.4b on the Antarctic Peninsula to date (https://scar.org/library-data/avian-flu, 08/04/2024). This region contains several important breeding areas for seabirds, as evidenced by the designation of multiple Antarctic Specially Protected Areas (ASPAs) and Important Bird Areas (IBAs), particularly in and around Marguerite Bay. This includes globally significant populations of Adélie Penguins *Pygoscelis adeliae*, South Polar Skuas, and Antarctic Shags *Leucocarbo bransfieldensis*, including populations that have been subject to long-term monitoring (BirdLife International 2025; Phillips et al. 2019).

Specifically, we aimed to detect past or ongoing mortality events driven by HPAI H5N1 2.3.4.4b and identify the species involved in Marguerite Bay and quantify the potential impact of mortality events on the species involved. Beyond gold-standard qPCR testing, we evaluated a rapid antigen/lateral flow test in a remote field settingallowing for rapid in field diagnostics, with potential for rapid notification and response.

## METHODS

### Study site

Fieldwork was undertaken on Dismal (68° 06’ S 68° 51’ W) and Horseshoe Islands (67° 50’ S, 67° 14’ W) in Marguerite Bay in the Western Antarctic Peninsula (Fig. 1). Dismal Island is the largest of the Faure Islands situated at the western end of Marguerite Bay, roughly 35 km south of Adelaide Island. It has an area of ~7 km^2^ and maximum elevation of 60 m and is the largest of 21 rocky islands/islets in a small archipelago (Karaoğlan et al. 2023). Horseshoe Island is ~70 km ENE of Dismal Island, with an area of 60 km^2^ and maximum elevation of 879 m, with significant peaks, glaciers, freshwater lakes, and other, diverse geomorphological features (Yıldırım 2019). There are four research stations in the region, of which only Rothera Research Station (United Kingdom) and San Martín Base (Argentina) are active year-round (Fig. 1). Horseshoe hosts an Antarctic Specially Protected Area (ASPA 181 – Farrier Col; ATS 2025), which protects the scientific and environmental values associated with a series of freshwater lakes in the centre of the island. Horseshoe Island is also the site of the Turkish Scientific Research Camp. Little is documented on the vertebrate fauna of either island. Horseshoe is only ~3 km from Lagotellerie Island (ASPA 115) which hosts important populations of Adélie Penguin, South Polar Skua, and Antarctic Shags (ATS 2025).

**Figure 1:**
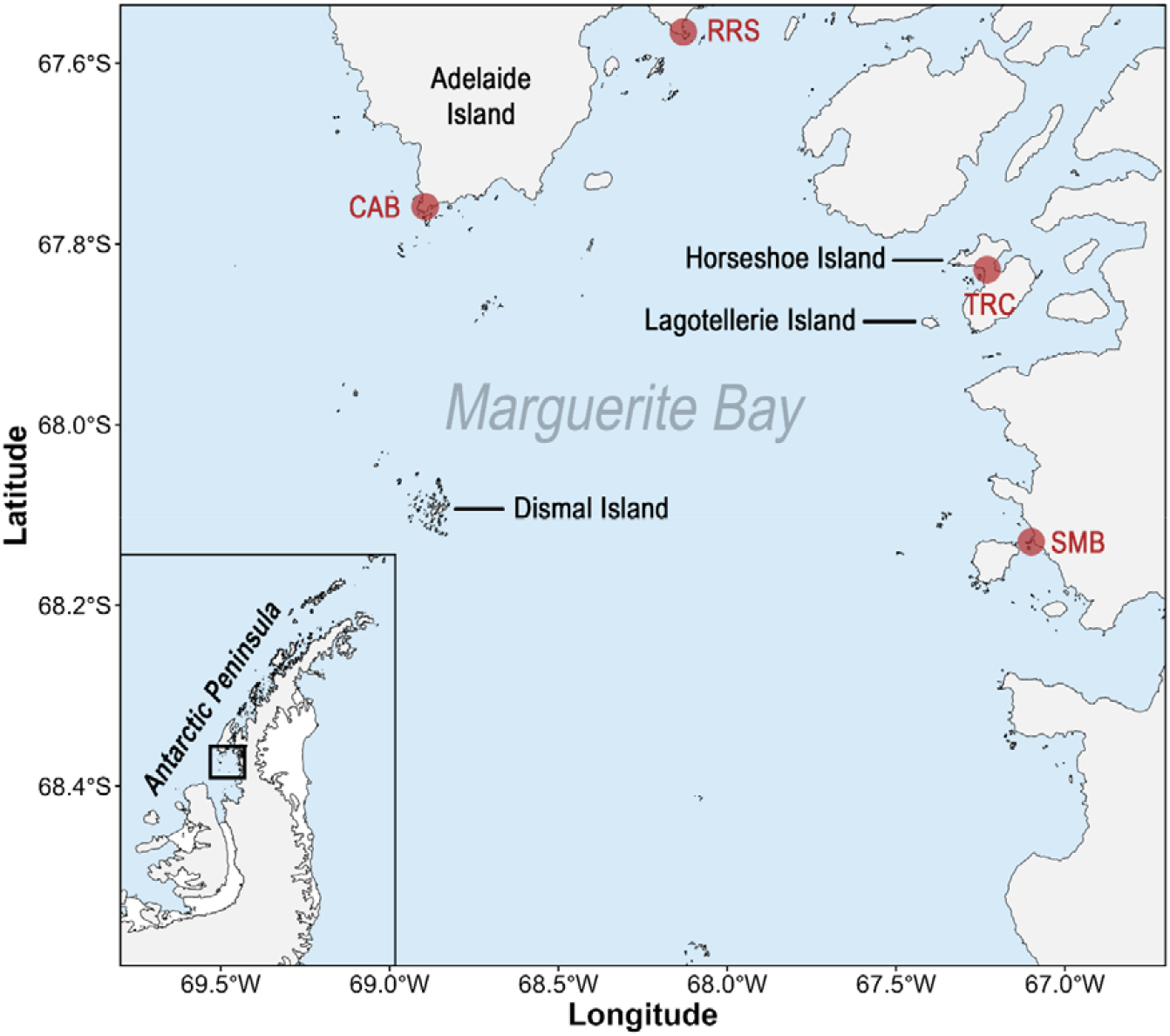
Marguerite Bay on the West Antarctic Peninsula (see inset), showing survey and sampling locations Dismal and Horseshoe Islands, nearby and geographically relevant Lagotellerie and Adelaide Islands, and the four research stations Carvajal Antarctic Base (CAB - Chile), Rothera Research Station (RRS – United Kingdom), Turkish Scientific Research Camp (TRC – Türkiye), and San Martin Base (SMB – Argentina).

### Field surveys and sampling

Fieldwork occurred during the 9^th^ Turkish Antarctic Expedition aboard the Betanzos Chilean research vessel between 16^th^ February and 5^th^ March 2025. During this period, fieldwork on Dismal Island occurred only on 16^th^ February (full day) and 5^th^ March (1.5 hrs) whereas fieldwork on Horseshoe Island occurred on most days between 17^th^ February – 5^th^ March.

Due to time constraints and difficulty navigating substantial ice and snow coverage, only the rocky region on the eastern side of the main northern bay of Dismal Island was surveyed for breeding seabirds and signs of HPAI (Fig. 1). Additionally, a Zodiac boat was taken around the island to observe breeding seabirds and marine mammals around the island, and to land on and survey two other adjacent islets on the 16th of February. Horseshoe Island was more comprehensively surveyed, particularly the central and northern sections, while the southern section was largely inaccessible due to challenging topography and glaciers (Fig. 1).

Live birds and mammals were recorded as part of a separate comprehensive ecological survey, of which some observations are used in this study (Table 1 – note scientific names are provided in this table, rather than in-text hereafter). Species, age, count, notes on injury/illness (e.g., lethargy, bloodshot eyes in birds, neurological symptoms), and location were recorded for every live bird. For South Polar Skuas only, a concerted effort was made to count the number of breeding pairs in surveyed areas, and the location of recorded nests/territories was used avoid double counting. Observations were carried out by 1-2 observers using binoculars, and if required, a digital camera. From these data, we quantified all species present, occurrence of breeding records, and suspect cases of HPAI in live birds and mammals. Carcasses classified as ‘fresh’ (eyeball present), ‘older’ (eyeball absent, otherwise no clear sign of decay), or ‘decayed’ (carcass in state of decay). Whether the carcass had any sign of having been scavenged was also recorded. Only carcasses deemed ‘suspect’ for HPAI H5N1 2.3.4.4b (including those which subsequently tested positive) were included. This required the carcass to be of the whole body (e.g., not just wings left from a predation/scavenging event), and to be in a condition that facilitated sampling of the cloaca, oropharynx, or brain (i.e., not severely decayed such that these samples could not be collected if the carcass was selected for sampling). Additional observations of behaviours or situations which may facilitate viral transmission were documented.

**Table 1:**
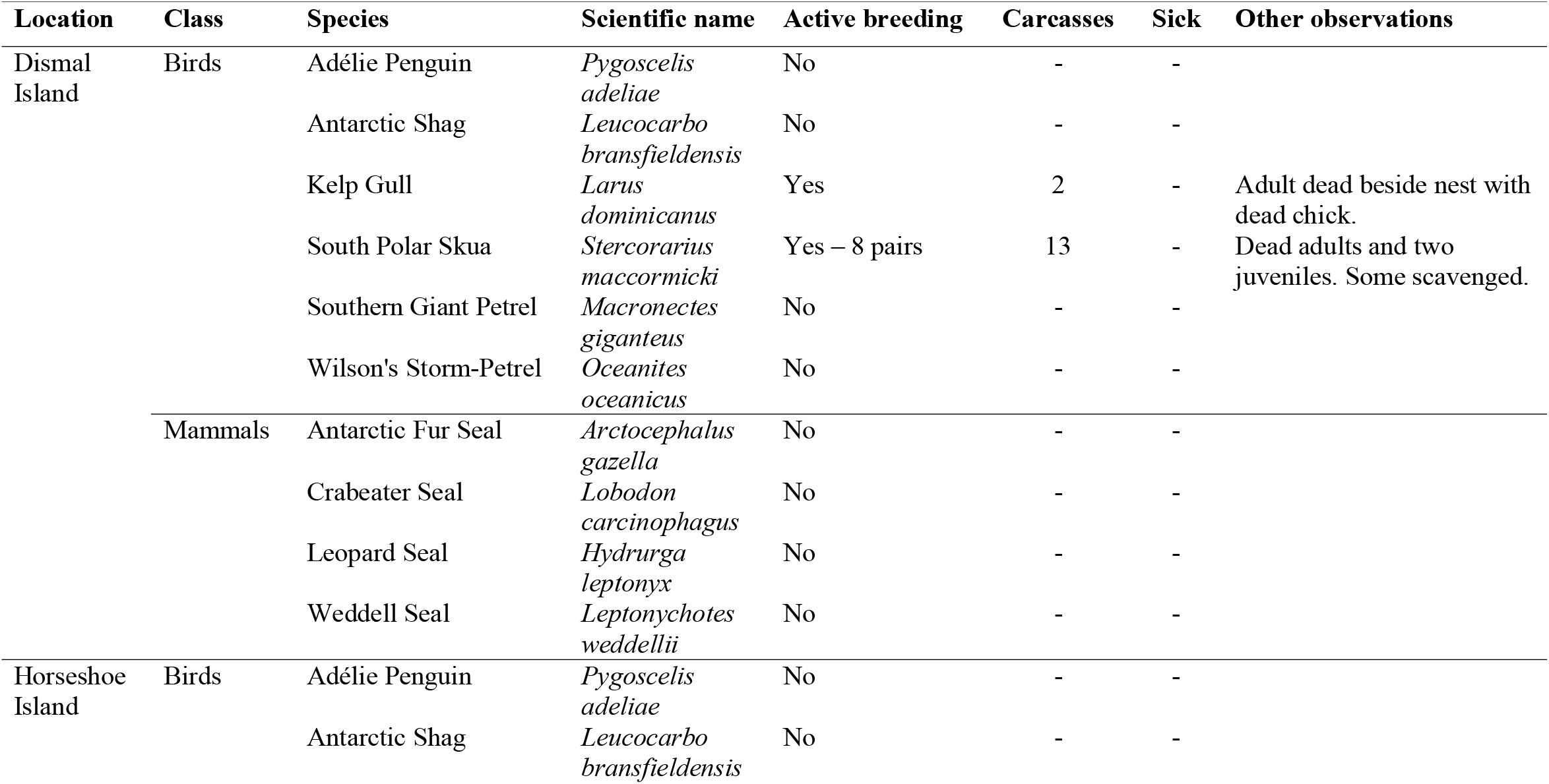
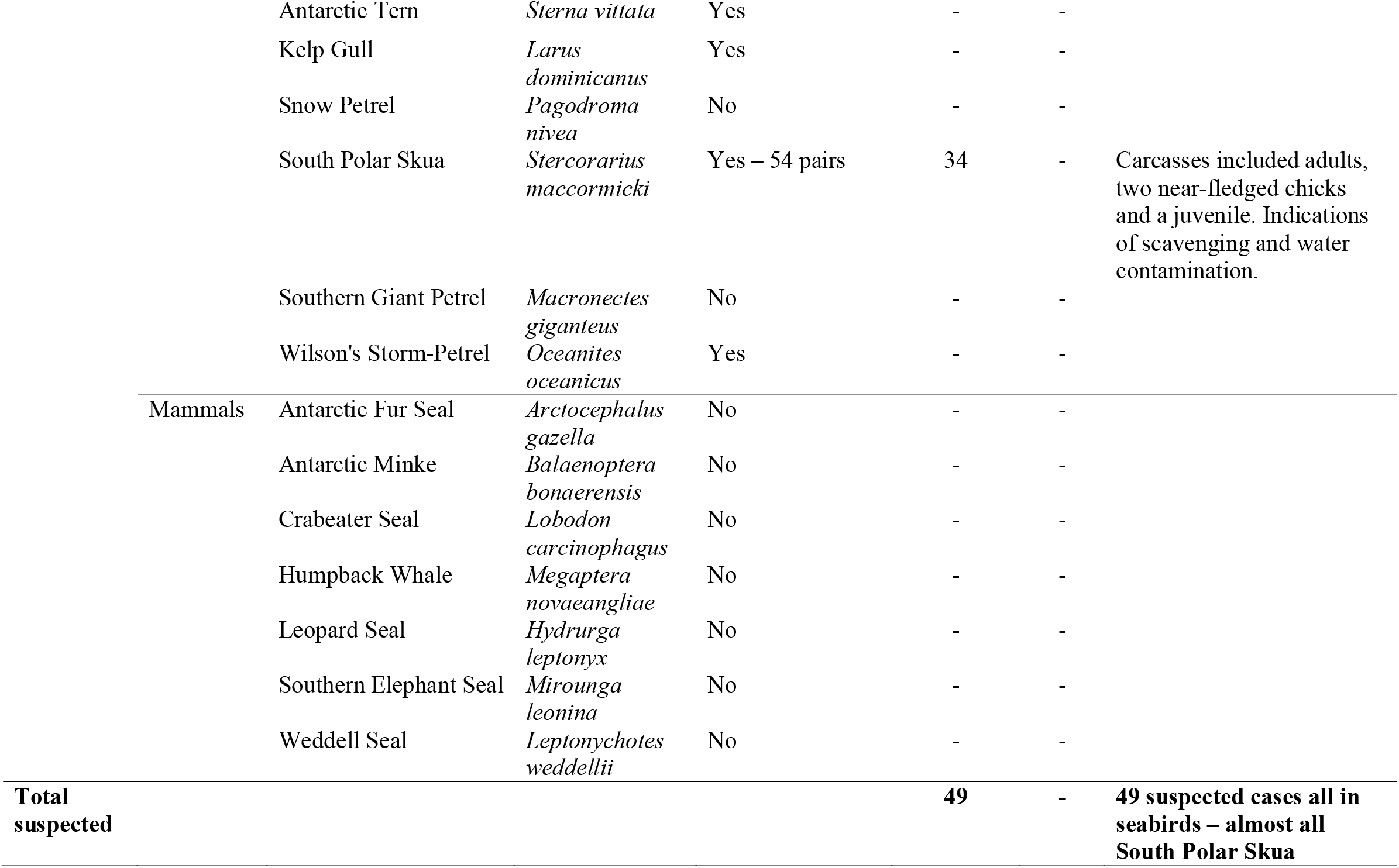
Birds and mammals from Dismal and Horseshoe Islands, Marguerite Bay from 16^th^ February – 5^th^ March 2025 with records of active breeding (quantified for South Polar Skua), counts of suspected HPAI H5N1 2.3.4.4b carcasses and sick individuals, and other relevant observations.

Only seabirds were sampled for HPAI, which involved collecting cloacal and oropharyngeal swabs, brain swabs, or swabs of fresh faeces.All sampling was done in accordance guidance and protocols from COMNAP, SCAR, and the 9^th^ Turkish Antarctic Expedition.

### Influenza A virus detection and H5 confirmation

In the field, one swab was placed in the dilution buffer provided with the VDRG® AIV Ag Rapid kit 2.0 (Median Diagnostics), after which four drops of the solution were placed on the sample device (as per instructions: https://www.mediandiagnostics.com/en/21_view?idx=26). This test kit detects both, low and high pathogenicity avian influenza viruses (AIVs), meaning a true positive indicates the presence of AIVs but not their type. The result was read at the recommended 15-minute time interval but were kept for at least 3 hours and read at 40 minutes and up to 3 hours. Given that the ambient temperature during sampling in the region was generally below 0°C, and the minimum recommended storage temperature of the test is 2°C, we suspect that the colder ambient conditions prolonged the time to a positive result. Two further swabs were collected from each bird and stored in universal transport medium (Macrogen) at room temperature for up to 17 days (depending on sampling date). These were subsequently stored at 4°C for five days, after which they were deposited in a denaturant viral transport media (KaiBiLi™ Extended NTM, The Hague, Netherlands). From these swabs, RNA was extracted using the ZYBIO PRT DNA & RNA Extraction Kit in a ZYBIO Nucleic Acid Isolation System EXM3000. RNA was assayed by reverse transcriptase real time PCR for detection of influenza A virus targeting the matrix (M) gene (WHO 2021). Results with a cycle threshold (Ct) <35 were considered positive, greater than 35 were considered inconclusive, and those with CT values greater than 40 were deemed negative. To confirm the presence of HPAIV H5N1 clade 2.3.4.4 from matrix positive RNA, were further analyzed using three additional real-time RT-PCR assays for H5, the HA multibasic cleavage site, and NA genes in accordance with VSL-USDA protocols (NVSL protocols 1732.02, 1767.01, and 1768.01) (as performed in Ulloa et al. 2023; Bennett-Laso et al. 2024).

## RESULTS

### Field surveys

Across both islands, 15 bird and mammal species were recorded as either breeding or present (10 on Dismal, 15 on Horseshoe). Active breeding was observed in Kelp Gull and South Polar Skua on Dismal Island, and in both species as well as Antarctic Tern and Wilson’s Storm-Petrel on Horseshoe Island (Table 1). For South Polar Skua, at least eight pairs were actively breeding on Dismal Island, and at least 54 pairs were actively breeding on Horseshoe Island. Across the entire survey period and both locations, 49 carcasses were found, with 15 on Dismal Island (one Kelp Gull adult and chick and 13 South Polar Skua), and 34 on Horseshoe Island (all South Polar Skua) (Table 1). Only one carcass (on Horseshoe) was fresh; all others were older, and none were decayed. Carcasses were typically found in rocky areas close to breeding areas and freshwater sources, often where healthy breeding adults and young were also present. We did not find any fresh or older mammal carcasses and did not observe any live birds or mammals with disease signs.

Through observations, we aimed to identify putative transmission pathways for HPAI. Specifically, we observed groups of typically 2-10 South Polar Skuas bathing and drinking from Skua Lake in the northern section (during three visits) and the freshwater lakes within the ASPA 181 area (during six visits) (Fig. 3a). Over six visits, we observed 10-30 Antarctic Terns of all ages bathing and drinking from one of the lakes within ASPA 181 which the skuas were also using (Fig. 3b). During one cold day when Skua Lake was almost completely frozen, a group of 49 skuas was observed bathing, drinking, and engaging in social behaviour in a confined area (Fig. 3c). Five skua carcasses were observed partially submerged on the edge of Skua Lake and we also observed skuas defecating into the lake on five occasions, although given their ongoing presence at this site, this behaviour must be commonplace (Fig. 3d-e). Active scavenging was not observed, however many carcasses of South Polar Skuas had been partially eaten, likely due to cannibalistic predation and/or scavenging (n=1 on Dismal, n=8 on Horseshoe), and when tested, these eaten carcasses (n=3) were positive for HPAI H5N1 2.3.4.4b (Fig. 3f). Some carcasses were found 1-5 m from Adélie Penguins (n=7) and Antarctic Fur Seals (n=1). While neither species was observed to interact with the carcasses, Adélie Penguins frequently ran their bills through and ate snow, which if contaminated could facilitate spread. Around South Polar Skua nests, anecdotal observations of bird carcasses which had been eaten included Kelp Gulls, other South Polar Skuas, Antarctic Terns, and Snow Petrels.

### Confirmed HPAI H5N1 2.3.4.4b cases

A total of 30 samples were collected: 22 from South Polar Skua carcasses (n=6 on Dismal Island, n=16 on Horseshoe Island), seven faecal samples from alive and apparently healthy South Polar Skuas (n=1 on Dismal Island, n=6 on Horseshoe Island), and 1 from a Kelp Gull carcass on Dismal Island (Table 2). Of these 30 samples, 20 were positive using in-field rapid antigen tests (RATs), and 23 were confirmed by real time RT-PCR (Table 2). There was high agreement (90%) between the results from RATs and real time RT-PCR analysis (Table 2). RATs produced no false positive results. Three false negatives were detected (thus, provisionally, a 10% false negative rate) – two from brain swabs of a Kelp Gull (Ct=30.10) and a South Polar Skua (Ct=22.40), and one from a cloacal/oropharyngeal swab of a South Polar Skua (Ct=35.69) (Table 2; Appendix A1). Cycle threshold (Ct) values ranged from 18.91 – 35.69, with a median value of 27.39 (Appendix A1). As all carcasses by 1 were in the “older” category, we were unable to address whether carcass age has an impact on RAT performance.

**Table 2:** Sampling effort for HPAI H5N1 2.3.4.4b from Dismal and Horseshoe Islands, Marguerite Bay, Western Antarctic Peninsula, summarised by species and swab type, results from rapid antigen and M-gene PCR contrasted by false positives and negatives.

| Location | Species | Swab sample type | Number of samples | Positive RAT | Positive M-gene PCR | RAT false positive | RAT false negative |
| --- | --- | --- | --- | --- | --- | --- | --- |
| Dismal Island | South Polar Skua | Brain | 1 | 1 | 1 | - | - |
|  |  | Cloacal/Oropharyngeal | 5 | 4 | 5 | - | 1 |
|  |  | Scat | 1 | - | - | - | - |
|  | Kelp Gull | Brain | 1 | - | 1 | - | 1 |
| Horseshoe Island | South Polar Skua | Brain | 10 | 10 | 10 | - | - |
|  |  | Cloacal/Oropharyngeal | 6 | 5 | 6 | - | 1 |
|  |  | Scat | 6 | - | - | - | - |
| Total |  |  | 30 | 20 | 23 | 0 | 3 |

HPAIV H5 clade 2.3.4.4, was confirmed by real time PCR (both subtype H5 clade 2.3.4.4 and pathotyping assays), and all positive samples reported similar Ct Values as gen M. In contrast, none of the samples reported Ct value for N1 subtyping. To complement this result, N1 amplicons were ran in a 2% electrophoresis observing bands at similar size as N1 positive control.

## DISCUSSION

HPAI outbreaks are occurring across Antarctic and subantarctic regions as HPAI H5N1 2.3.4.4b spreads, underscoring the need for ongoing assessment of species’ susceptibility, monitoring of transmission dynamics, and enhanced detection capabilities (Banyard et al. 2024). At the southern extent of known spread along the Western Antarctic Peninsula to date, we detected HPAI H5N1 2.3.4.4b mortalities in South Polar Skuas on distinct islands in Marguerite Bay, as well as one confirmed and one suspected case Kelp Gulls, and no other signs of the disease in any other seabirds or mammals at the time of sampling. The initial viral detection was done using rapid antigen tests which successfully detected AIV presence/absence in 90% of cases, which was confirmed using real time RT-PCR. However, despite observations of many situations which could facilitate interspecific spread, we found limited evidence of widespread impact on other seabirds or mammals. Disproportionate mortalities in South Polar Skua align with global findings on susceptibility of skuas for HPAI (Camphuysen et al. 2022; Gorta et al. 2024), providing valuable insights into the broader impacts of these outbreaks in vulnerable Antarctic ecosystems. Our findings enhance our understanding of the broader ecological consequences of HPAI H5N1 2.3.4.4b outbreaks and will benefit ongoing monitoring efforts, as well as conservation and biosecurity measures in the context of the ongoing spread.

South Polar Skuas were overrepresented in our observations of suspected and confirmed HPAI mortalities, making up 21 of the 22 tested carcasses, and 47 of the 49 suspect carcasses found during surveys. Of note is that Brown Skuas, another species overrepresented in HPAI cases in the Antarctic, are not present in Marguerite Bay. The susceptibility of skuas to HPAI H5N1 2.3.4.4b is well-documented on a global scale (Camphuysen et al. 2022; Gorta et al. 2024), although to our knowledge this is the first detailed examination of an outbreak in South Polar Skuas. Mortalities were concentrated around breeding areas and freshwater sites, where social behaviours, cannibalistic predation and scavenging, and contaminated freshwater sources (faeces and submerged carcasses) likely facilitated viral spread, as in other skua species (Fig. 3; Banyard et al. 2022; Camphuysen et al. 2022; Gorta et al. 2024). Only one carcass was fresh with intact eyes, while the rest were older and without eyes, suggesting the bulk of mortalities had occurred prior to sampling, and that limited mortality was occurring during the sampling period. This was additionally supported by no new, fresh carcasses, or live birds with signs of disease, being observed in the central Horseshoe or northern Horseshoe regions where repeat sampling occurred (see Fig. 2 insets 2 and 3). While we detected 34 South Polar Skua carcasses on Horseshoe Island where comprehensive surveys for breeding pairs were undertaken, 54 actively breeding pairs were also observed, primarily feeding mature chicks. Although no baseline exists for South Polar Skua breeding success on Horseshoe Island – a metric that varies annually in the region (Phillips et al. 2019) – our observations show that despite widespread mortality, many individuals were still breeding successfully at the time of surveys late in the breeding season.

**Figure 2:**
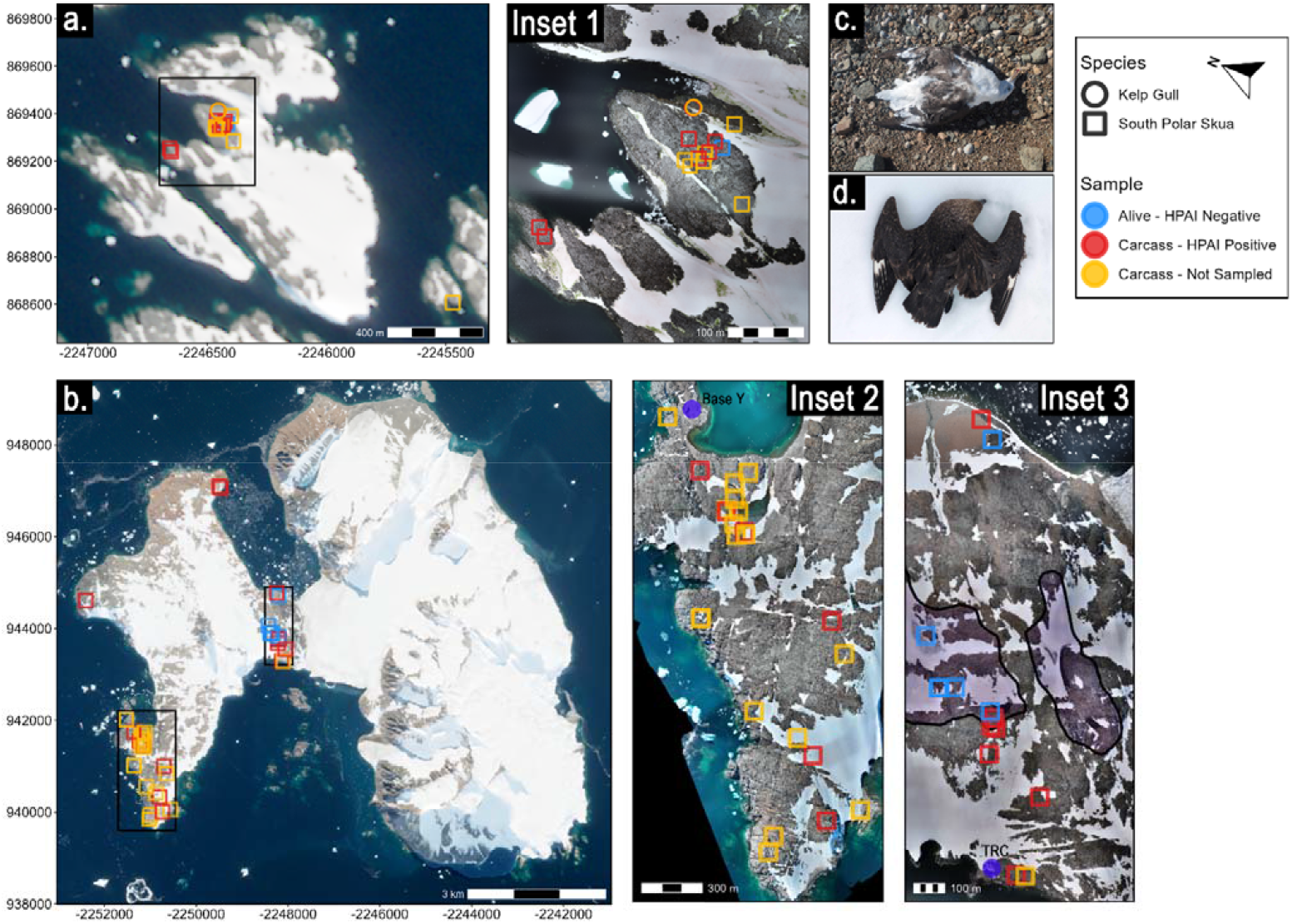
Locations of all samples and suspected HPAI H5N1 2.3.4.4b carcasses on Dismal (**a**.) and Horseshoe Islands (**b**.) with detailed locations shown in three insets (whole island imagery sourced from Sentinel-2; high resolution inset imagery provided by TÜBİTAK; north arrow in the legend applies to all maps in projected coordinate system WGS 84/Antarctic Polar Stereographic - EPSG:3031). Locations of Base Y and the Turkish Scientific Research Camp (TRC) can be found in insets 2 and 3 respectively (labelled purple points) and the area of Farrier Col ASPA 181 is highlighted in light purple in inset 3. Additional images are of positive HPAI carcasses of an adult Kelp Gull *Larus dominicanus* (**c**.) and a South Polar Skua *Stercorarius maccormicki* (**d**.).

**Figure 3:**
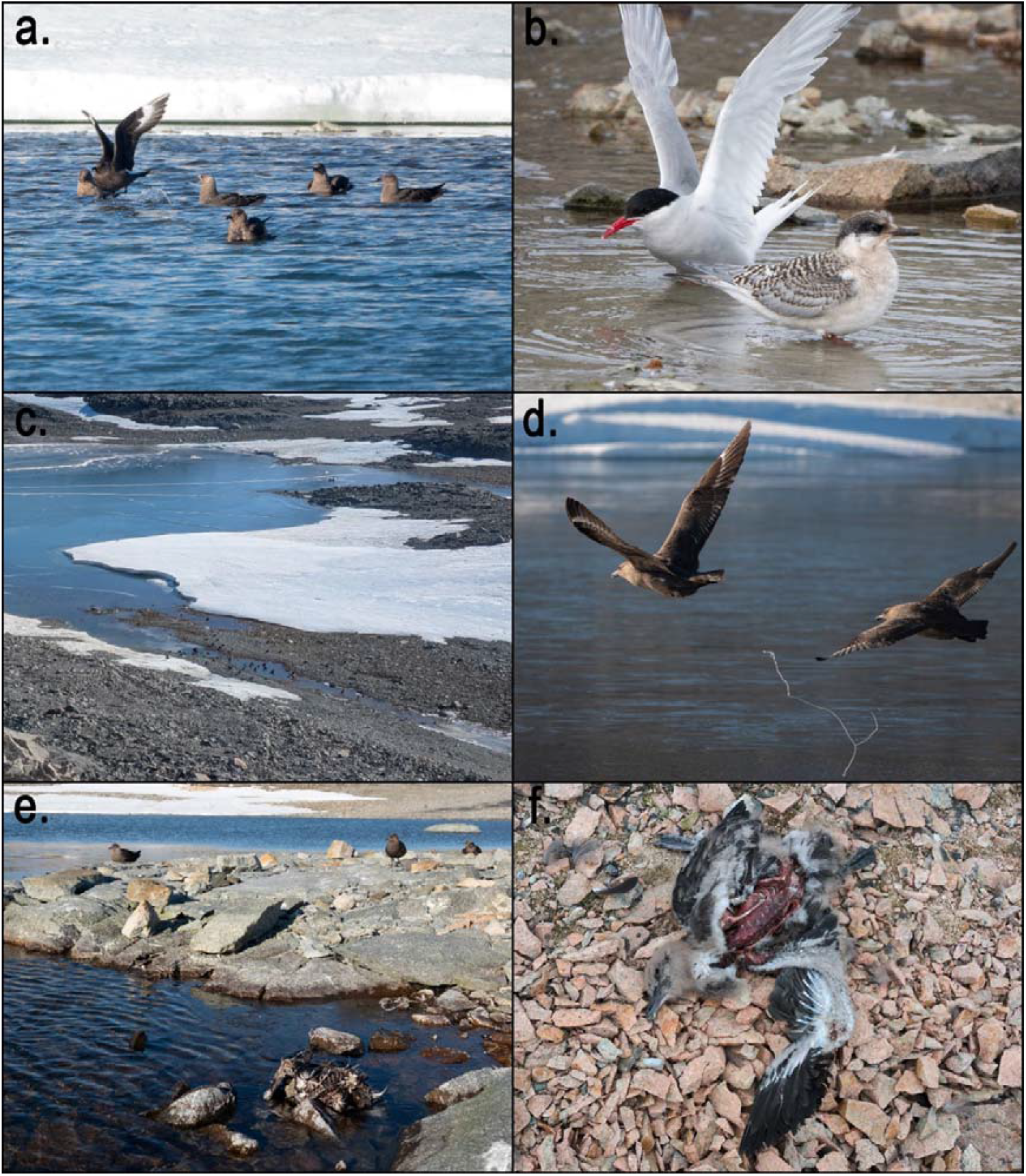
Behavioural and situational factors that may facilitate spread of HPAI H5N1 2.3.4.4b included: communal drinking and bathing in freshwater pools and lakes in South Polar Skua *Stercorarius maccormicki* (**a**.) and Antarctic Tern *Sterna vittata* (**b**.), the former sometimes congregating in large groups at these sites and engaging in social behaviours (**c**.). South Polar Skua were also observed to defecate in communal freshwater areas (**d**.) and multiple carcasses were found partially submerged in Skua Lake (**e**.). Scavenging of young and adult carcasses was common (**f**.). Photographs by S. Gorta.

Aside from mortalities in South Polar Skuas and two Kelp Gulls, mortalities and signs of HPAI infection in live birds and mammals were not detected. The Kelp Gull carcasses were an adult and presumably its chick on Dismal Island near the mass mortality of South Polar Skuas (Fig. 2). Kelp Gulls are opportunistic scavengers which could include skua carcasses (Favero et al. 1997) and may also have been exposed to contaminated meltwater at the site, either of which may have led to infection. No pinnipeds, including those in a large mixed haul-out site of four species (Antarctic Fur Seal, Crabeater Seal, Southern Elephant Seal, and Weddell Seal) on the north-western point of Horseshoe Island showed any indications of past or current HPAI infection, despite one skua carcass being found within the haul-out site. The same applied for large moulting groups of Adélie Penguins on Horseshoe Island, some of which were <3 m away from positive HPAI skua carcasses, and frequently were observed eating snow, but none showed any signs of infection. This does not rule out that some penguins or pinnipeds, may have been asymptomatic or presented subtle signs which went undetected. Additionally, Antarctic Terns were observed breeding next to, drinking from, and bathing in freshwater lakes within the ASPA 181 region on Horseshoe Island, where South Polar Skuas also frequently drank and bathed and positive HPAI carcasses were found (Fig. 2; Fig. 3b). Despite this, no live or dead Antarctic Terns were found that were suspected to be HPAI positive. Elsewhere, terns have been subject to mass mortality events, and in some instances, population declines in globally significant populations (Rijks et al. 2022; Pohlmann et al. 2023; Uhart et al. 2024). Compared to areas where terns have been severely affected by HPAI, the relatively low breeding densities and limited interspecific close contact, and high-wind conditions at Horseshoe Island likely reduced transmission risk. As such, although we did not detect any evidence of HPAI in Antarctic Terns, contamination of communal freshwater sites represents a plausible infection pathway for this species in Antarctica and should be monitored in ongoing surveillance efforts.

Ongoing surveillance of important and vulnerable bird and mammal populations is paramount to managing outbreaks in known areas and preparing for and detecting outbreaks in novel regions. Real time PCR detection and confirmation remains the gold standard for HPAI, with a multistep approach imperative to confirm not only H5, but the clade (2.3.4.4b) and pathotype. Notably, the N1 real-time PCR assay yielded negative results; however, electrophoresis of the PCR products showed patterns compatible with N1, confirming HPAI H5N5 and strongly suggesting that the TaqMan-based probe may require updating. Similar lack of N1 detection has been observed in other regions in Antarctica this season (https://scar.org/library-data/avian-flu).

Due to the remoteness of Marguerite Bay (and Antarctica in general), we employed a rapid test approach, and successfully detected avian influenza virus which was verified subsequently to be HPAI H5N1 2.3.4.4b in South Polar Skua carcasses using VDRG® AIV Ag Rapid kit 2.0 (Median Diagnostics) with a 10% false-negative rate. Given the advantages of rapid tests, we recommend the application of these tests in a field setting in three situations: (1) when no other testing kits are available; (2) when results are needed rapidly and no other rapid testing approaches are possible; and (3) to boost sampling and detection effort with limited available testing kits. However, these tests have a number of shortcomings, and there is need for further evaluation. Critically, interpretation of negative results should be informed by in-field observations and context, given their potentially reduced sensitivity which may lead to false negatives. Carcass age should also be considered: despite our ability to detect HPAI in older carcasses, it is unclear whether decomposition or sample type would have an impact on the detection performance. The impact of temperature should also be considered, as outdoor conditions in Antarctica are outside the range of approved temperatures for the use of rapid tests. Indeed, we found that the 15 min read window was not always appropriate. Importantly, these tests did not distinguish between high and low pathogenicity avian influenza, which has serious implications for interpretation and subsequent actions and management (Wille et al. 2024). Finally, while the VDRG® AIV Ag Rapid kit 2.0 worked well in this context, different rapid tests may have considerably difference performance and should be evaluated for HPAI H5 detection prior to use. In practice, with careful consideration of their limitations, the context of the environment of use and the species on which they are used, these are an effective tool for detecting avian influenza viruses during HPAI outbreaks in wildlife and should be incorporated into the surveillance toolkit.

This toolkit would further benefit from cross-disciplinary collaboration to better understand the links between HPAI outbreaks and real ecological outcomes. Long-term population monitoring complemented by increased efforts to understand the year-round movements of Antarctic seabirds and mammals for which pre-HPAI baselines are known (e.g., Phillips et al. 2019) will be essential for assessing the full impact of HPAI H5N1 2.3.4.4b on Antarctic wildlife populations and informing conservation strategies. This call for greater collaboration extends to sampling efforts as well, emphasising the need to maximise limited conservation and research resources while fostering cooperation and reducing redundancy. Further sampling efforts in Marguerite Bay would build on our findings by increasing the number of live individuals sampled, and both live and dead individuals from species that are common, potentially vulnerable, or of conservation concern (e.g., Antarctic Fur Seal, Antarctic Tern, Adélie Penguin). Additionally, genomic sequencing could provide valuable insights into the introduction of HPAI to Marguerite Bay, helping to clarify its potential origins and pathways in the broader context of Antarctic and subantarctic spread (e.g., Clessin et al. 2025).

HPAI H5N1 2.3.4.4b poses a global conservation risk to ecosystems, populations, and species in Antarctica, with the added uncertainty around its ecological impact associated with continued eastward spread into regions. We surveyed seabirds and mammals at the frontier of HPAI’s southward spread in Marguerite Bay, detecting mortalities in South Polar Skuas (many still breeding successfully) and limited mortality in other species. The ongoing spread of the virus into new regions (Banyard et al. 2024; Clessin et al. 2025), combined with uncertainties regarding its long-term ecological impact, necessitates ongoing surveillance and proactive conservation measures. Continued long-term monitoring of populations will be central to understanding the population-level impacts of HPAI as it continues its spread throughout the world’s bird and mammal populations.

## ACKNOWLEDGEMENTS

This study was carried within the Ninth Turkish Antarctic Expedition (TAE-IX), under the auspices of the Presidency of the Republic of Türkiye, supported by the Ministry of Industry and Technology, and coordinated by TUBİTAK MAM Polar Research Institute. We thank Doğaç Baybars Işıler for his logistical support and assistance as part of the TAE-IX. We would like to acknowledge the Korea Polar Research Institute (KOPRI) for providing sampling kits, and Chilean Antarctic Institute (INACH), particularly Marcelo González-Aravena, for facilitating inactivation of samples. ÇH□ thanks H. Batubay Özkan and Barbara Watkins for their support of the Biodiversity and Conservation Ecology Lab at the University of Utah School of Biological Sciences. This study was partially funded by INACH Regular RT-0821 and Fondecyt 1211517. The WHO Collaborating Centre for Reference and Research for Influenza is supported by the Australian Department of Health.

